# Constraints and trade-offs shape the evolution of T cell search strategies

**DOI:** 10.1101/2022.07.28.501835

**Authors:** Inge M N Wortel, Johannes Textor

## Abstract

Two decades of *in vivo* imaging have revealed how diverse the shapes and motion patterns of migrating T cells can be. This finding has sparked the notion of “search strategies”: T cells may have evolved ways to search for antigen efficiently and might even adapt their motion to the task at hand. Mathematical models have indeed confirmed that observed T-cell migration patterns resemble a theoretical optimum in several contexts; for example, frequent turning, stop-and-go motion, or alternating short and long motile runs have all been interpreted as deliberately tuned behaviours, optimising the cell’s chance of finding antigen. But the same behaviours could also arise simply because T cells *can’t* follow a straight, regular path through the tight spaces they navigate. Even if T cells can be shown to follow a theoretically optimal pattern, the question remains: has that pattern truly been evolved for this particular searching task, or does it merely reflect how the cell’s migration machinery and surroundings constrain motion paths?

We here examine to what extent cells can evolve search strategies when faced with realistic constraints. Using a cellular Potts model (CPM), where motion arises from interactions between intracellular dynamics, cell shape, and a constraining environment, we simulate an evolutionary process in which cells “optimise” a simple task: explore as much area as possible. We find that cells evolve several motility characteristics previously attributed to search optimisation, even though these features were *not* beneficial for the task given here. Our results stress that “optimal” search strategies do not always stem from evolutionary adaptation: instead, they may be the inevitable side effects of interactions between cell shape, intracellular actin dynamics, and the diverse environments T cells face *in vivo*.

## Introduction

T cells have the rare ability to migrate in nearly all tissues within the human body. In lymphoid organs, such as the thymus and lymph nodes, T cells must migrate to develop and get activated; in peripheral “barrier” tissues, like the lung, the gut, and the skin, T cells continuously patrol in search of foreign invaders. Although T cells stay motile in these different contexts, they do adapt their morphology and migratory behaviour to environmental cues. Naive T cells rapidly crawl along a network of stromal cells in the lymph node, alternating between short intervals of persistent movement and random changes in direction [1, 2, 3, 4]. This “stop-and-go” behaviour lets them cover large areas of the lymph node quickly, and seems to be a good strategy for finding rare antigens without prior information on their location [5, 6, 7, 8]. Developing T cells adopt a similar strategy to find their specific ligand during negative selection in the thymic medulla [9, 10]. By contrast, positive selection in the thymic cortex involves migration at much lower speeds—perhaps due to the broader distribution of positively selecting ligands in the thymus [11]. This remarkably flexible behaviour has been suggested to reflect different “search strategies”, whereby T cells maximise their chance of finding antigen [5]. ^1^

The idea of search strategies has interesting implications. If T-cell migration patterns are indeed optimised for some specific function (or several functions depending on context and environment), then comparing their “search efficiency” can help us make sense of how T-cell function relates to these diverse migratory behaviours [14]. However, two major problems currently limit the conclusions we can draw from this work.

First, such optimality reasoning hinges on a tacit but crucial assumption: that we observe a given behaviour because it has been selected through evolution. But in reality, it is far from certain that evolution can optimise migration at all [15]. The complex mapping between *genotype* (the genes controlling cell migration, which can be transferred to the next generation) and the resulting *phenotype* (the migratory behaviour we see) means that not all search strategies are evolvable through natural selection.

For example, the Lévy foraging hypothesis states that searchers can maximise their efficiency by carefully tuning their directional persistence, alternating between many short and a few long, high-speed runs. Yet cells do not have individual “speed” or “persistence” genes, and may not be able to evolve the one without affecting the other. In fact, a universal coupling between speed and persistence (UCSP) has been described for many migrating cells: faster cells move more persistently [16, 17, 18]. Thus, the cell’s migration machinery already poses constraints on the motion patterns cells can adopt. These constraints are strengthened further by the complex, crowded environment T cells typically migrate in; both *in vivo* imaging and *in silico* modeling have highlighted how strongly these external cues can affect T-cell shapes and migration patterns [3, 19, 20]. These constraints mean that (evolving) T cells can only “choose” from a limited range of motion patterns, and that their behaviour is more likely to reflect some kind of compromise than a true “optimum”.

Second, to determine how “optimal” a migration pattern is, we must make assumptions; after all, even though it may be very useful for a searching T cell to be in two places at once or to move at the speed of light, we typically do not consider these options in a search for optimal behaviours. Put simply: we can only assess the “efficiency” of a strategy *relative* to a set of other strategies we think the cell can adopt [15, 21]. Studies investigating T-cell search mostly use (variations of) random walk models for this purpose [5, 22, 23, 24, 25]. These fairly simple mathematical or agent-based models can produce different motility patterns depending on parameters, which directly reflect properties like cell speed and turning behaviour. For a given dataset, fitting these parameters yields a model of the “observed” strategy whose search performance we can assess on imaginary targets *in silico*. Thus, we learn whether the observed motion pattern was a good strategy for some searching task.

Such models, however, are hard to interpret. Model selection is difficult because the same data can often be explained by multiple models depending on exactly how migration is quantified [26, 25, 27]: for example, while Harris *et al*. have claimed that T cells in the brain follow Lévy flights to find rare pathogens [22], others [28] recently cautioned that similar statistics may arise through other mechanisms. Furthermore, the search efficiency found in such models can again strongly depend on the structure of the environment [29]—and even models that differ only slightly can still make very different predictions of the area cells can explore on larger time scales [30]. But most importantly, even if these models indeed show that a behaviour benefits some T-cell function, they cannot tell us whether the same behaviour could also have arisen for another reason altogether.

To unravel which migratory patterns truly *are* optimised for search, it does therefore not suffice to construct a random walk model showing that they are beneficial in some context or other. Instead, there are other crucial points to consider—Is the proposed “optimal” strategy something a cell could realistically adopt or evolve, given the biophysical constraints of its internal migration mechanism and environment? Which migration pattern would these constraints impose on the cell if no evolutionary pressures existed? Do we *need* an evolutionary explanation for the pattern in question, or could it simply be a side effect of dynamic cell motion in a complex environment? Whereas several studies have tested whether an observed migration behaviour could theoretically be optimal for some T-cell function [22, 23, 24, 25], these additional questions have largely remained unanswered.

Here, we therefore examine which migration patterns emerge spontaneously from the cell’s migration machinery and/or the environment, asking to what extent T cells can still evolve or tune search strategies within those constraints. Since these questions are impossible to answer in random walk models—which lack descriptions of the cell’s machinery, shape, and environment—we instead turned to a cellular Potts model (CPM). Specifically, we use an existing model called the Act-CPM [31], in which migration arises from a machinery where cell shape, environment, and motility interact. We previously showed that this model naturally captures many of the constraints acting on a migrating T cell: it reproduces the UCSP, explains how cell shape dynamics limit possible migratory patterns, and can simulate (T–)cell migration in a realistic tissue environment [13]. We now use this model to simulate an evolutionary process where T cells optimise a simple task: exploring as much area as possible. We find that T-cell migration behaviours previously interpreted as optimal search strategies can also emerge spontaneously *without* being beneficial for the task at hand.

## Model

In a CPM [32, 33], cells are dynamic pixel collections that move via so-called “copy attempts”: by copying their identity, they can “steal” pixels from another cell at their borders (Figure 1A). These identity changes are attempted at random, but constrained by a set of rules that assign them an energetic cost Δ*H*. As Δ*H* determines the success probability of each change, these energy rules ultimately govern cell behaviour in the model. For example, they can constrain a cell’s size, shape, or interactions with neighbouring cells (Figure 1B). Importantly, since all cells “compete” for pixels on the grid through the same global energy, cells naturally interact with each other in CPMs of multicellular environments.

**Figure 1:**
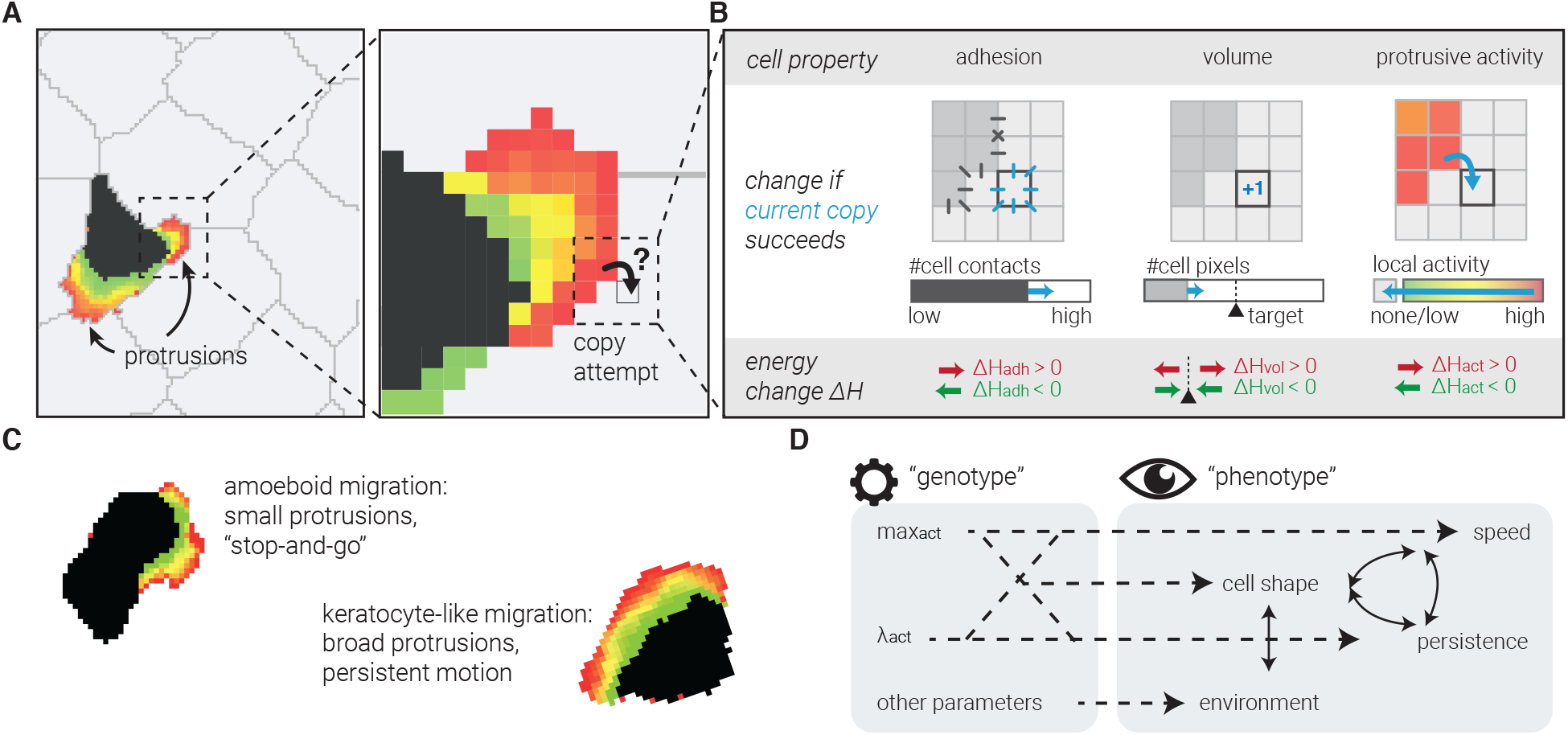
A computational model of cell migration with complex genotype-phenotype mapping. (A) CPM tissues are collections of pixels that each belong to one cell. Pixels try to copy their cell “identity” into neighbouring pixels of another cell. (B) Their success rate P_copy_ depends on how this would change the “energy” associated with different physical properties, such as surface tension (“adhesion”, left), or deviating from the normal cell size (“volume”, middle; an analogous constraint can be posed on the cell’s perimeter). The Act-CPM [31] adds another such property (right): each pixel’s “activity” represents the time since its most recent protrusive activity. Copy attempts from more into less active pixels are stimulated (negative ΔH_act_), placing positive feedback on protrusions. (C) Cells in the Act-CPM can have an amoeboid (stop-and-go) or a keratocyte-like (persistent) migration mode, which are associated with different cell shapes. (D) Complex genotype-phenotype mapping in the Act-CPM.

We have previously developed a CPM that models active cell migration based on actin dynamics [31, 13]. In the “Act-CPM”, pixels newly gained by a protruding cell retain their “protrusive activity” for some time. When these pixels then try to copy their own identity—extending the cell with yet another pixel—their activity makes them more likely to succeed: this positive feedback gives recently active pixels a better chance of protruding again (by assigning Δ*H*_act_ < 0; Figure 1B). Two parameters control this feedback: max_act_ controls how long pixels retain their protrusive activity, while λ_act_ tunes the protrusive strength relative to the other forces acting on the cell (i.e. those in Figure 1B). Together, max_act_ and λ_act_ control cell shape and motility. This model not only simulates cells that actively move by forming protrusions, but also reproduces different migration modes with their own protrusion shapes and motility patterns (Figure 1C) [31]. Qualitatively, it resembles the stop-and-go motility characteristic for T cells in the lymph node [1, 4].

Importantly, we have previously shown that this model also reproduces the UCSP [13]. Speed and persistence *emerge* as outputs of an intrinsic migration mechanism acting in a complex environment, rather than being imposed by the user. In evolutionary terminology, we speak of a complex mapping from *genotype* (fixed, cell-intrinsic values of the max_act_ and λ_act_ parameters) to *phenotype* (migratory pattern), where the genotype does not control the phenotype in any direct or obvious way (Figure 1D). Instead, interactions between the cell-intrinsic migration machinery, the cell’s shape, and the structure of the surrounding tissue dynamically determine the speed and direction of motion. This essential property allowed us to examine to what extent “optimal” search behaviour could evolve in a system with such a non-trivial genotype-phenotype mapping.

## Results

### Act cells can evolve migratory behaviour in a simple evolutionary algorithm

We therefore simulated a simplified form of evolution by means of a genetic algorithm (Figure 2A, see Methods for details). We first let cells evolve to explore as much area as possible in an empty environment with no surrounding tissue (Interactive Simulation S1). While this environment is not representative of what T cells encounter *in vivo*, it allowed us to see which migratory patterns could evolve without any constraints from the tissue. Since the evolutionary objective (having a large fitness) requires cells to explore a large area (Figure 2A), the theoretical optimum in this scenario is simply to maximise both speed and persistence—so as to move as far as possible without turning and visiting the same area twice [34]. To see if this theoretical optimum could arise through evolution of max_act_ and λ_act_, we started with a population where both parameters were too low for active cell migration (Figure S1A) and allowed them to evolve. During the evolutionary run, the average values of max_act_ and λ_act_ in the population gradually increased before eventually plateauing at values of 50 and 1165, respectively (Figure 2B). Strikingly, the same stable endpoint was reached in 9 out of 10 independent runs (Figure 2B; the last run did not fully converge but nevertheless moved towards that same point). This end point was associated with the highest fitness (Figure 2C)—suggesting that this parameter combination was somehow optimal. These results demonstrate that some form of evolutionary adaptation is taking place.

**Figure 2:**
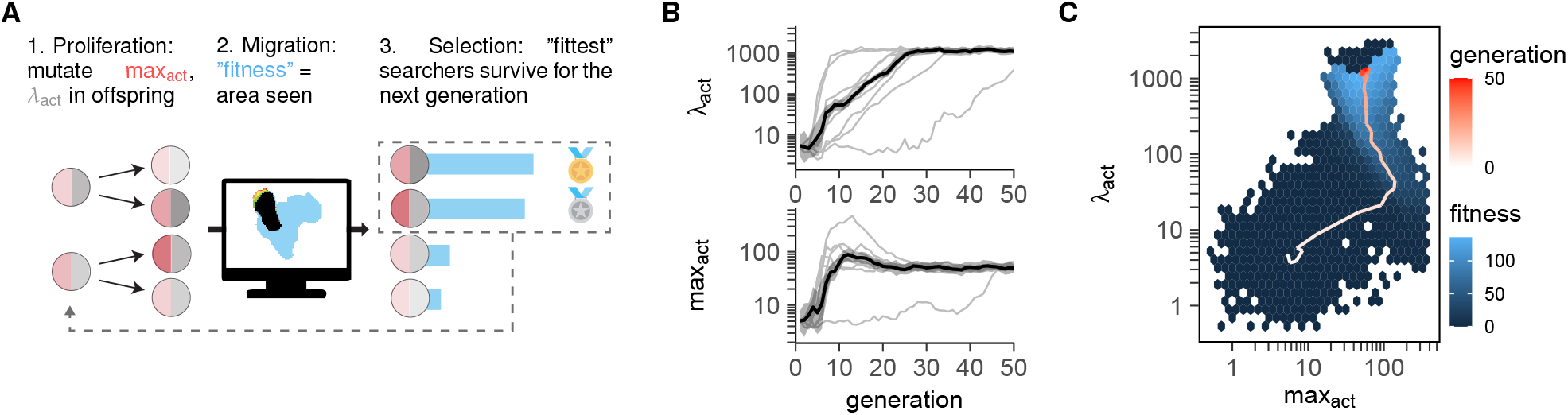
Evolution of optimal search behaviour in Act cells is subject to constraints and trade-offs. (A) Simulated evolution in a genetic algorithm. A population of 10 Act cells with their own (max_act_,λ_act_) parameters each produce three daughter cells with randomly “mutated” parameters (see Methods for details). After simulating migration for all 40 cells, only the 10 “fittest” cells (i.e. those that explored the largest area) survive as the next generation. (B) Evolution of λ_act_ and max_act_ over 50 generations. Black line + shaded area shows the mean ± SD within each generation. Thin gray lines show the same curve for 9 other, independent runs. (C) Evolution of λ_act_ and max_act_ in the context of the median “fitness” experienced by cells with those parameters. Fitness is defined as the area explored by the simulated cell (measured in number of cell areas of 500 pixels) and is zero if the cell breaks during the simulation. The red trajectory represents one single run; the (blue) fitness landscape is constructed by averaging measured fitnesses from all cells of all (10) independent runs at given parameters.

### Constraints and trade-offs limit evolution of optimal Act-cell search

To investigate how the evolved migratory behaviour arose, we next analysed motion at different parameter combinations along the evolutionary trajectory and surrounding the evolved optimum (Figure 3). The increase in the motility parameters max_act_ and λ_act_ coincided with an increase in migratory ability over the generations as measured by the average explored area (Figure 2C) as well as speed and persistence (Figure 3A). Along the trajectory, motion was well-described by a persistent random walk (Figure S2).

**Figure 3:**
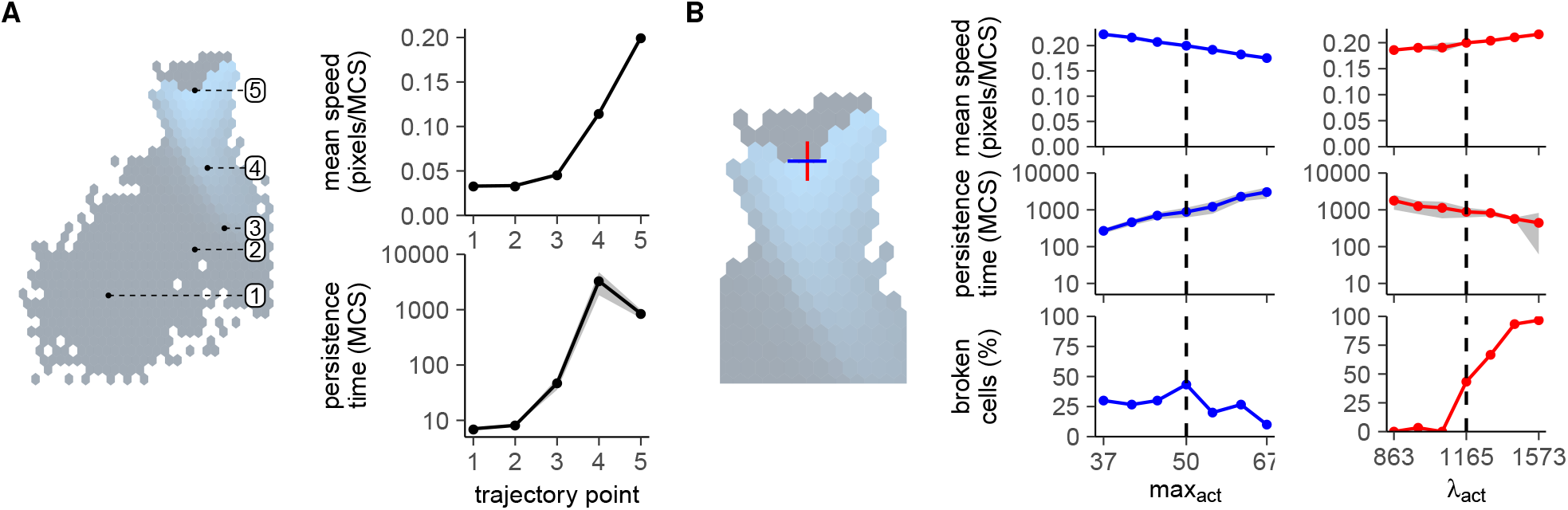
Evolution of optimal search in Act cells is subject to constraints and trade-offs. (A) Speed and persistence measured at different points in the fitness landscape of 2C. (B) Speed, persistence, and cell breaking measured around the evolved optimum (max_act_ = 50, λ_act_ = 1165).

Yet, intriguingly, cells at the empirical optimum did not have the theoretical maximum speed and persistence (Figure 3B). Whereas cells could still reach higher speeds by further increasing λ_act_, the higher force on the cell membrane also caused frequent cell breaking (Figure 3B). Thus, cells appear to evolve their λ_act_ to maximise speed while still conserving their integrity (Figure S1B). Likewise, there appears to be a trade-off between speed and persistence at parameters surrounding the optimum: the increased speeds observed at higher λ_act_ and lower max_act_ values come at the cost of a lower persistence (Figure 3B). This conflict likely arises because the evolved “optimal” cell is already quite broad and persistent (Movie S1). We have previously shown that, as the cell broadens, both speed and persistence saturate [13]. In this “saturation regime” of the UCSP, an even higher persistence requires a large effort to maintain a stable, broad protrusion, slowing the cell down [13]. Because of this trade-off, the cell evolves towards parameters where it is persistent enough that it rarely visits the same area twice (Movie S1), yet not so persistent that this comes at the cost of a low speed (Figure 3A,B). These results show that theoretical optima for speed and persistence may not be attainable when genotype-phenotype mapping is complex and gives rise to trade-offs.

### Environmental constraints, not evolved cell-intrinsic parameters, are the major determinants of migration patterns in tissues

Finally, we examined how the addition of environmental constraints affected the ability of cells to evolve optimal search behaviour. We therefore repeated the evolution experiment, but now assessed fitness by simulating T-cell migration inside a tissue instead of empty space (Figure 4A).

**Figure 4:**
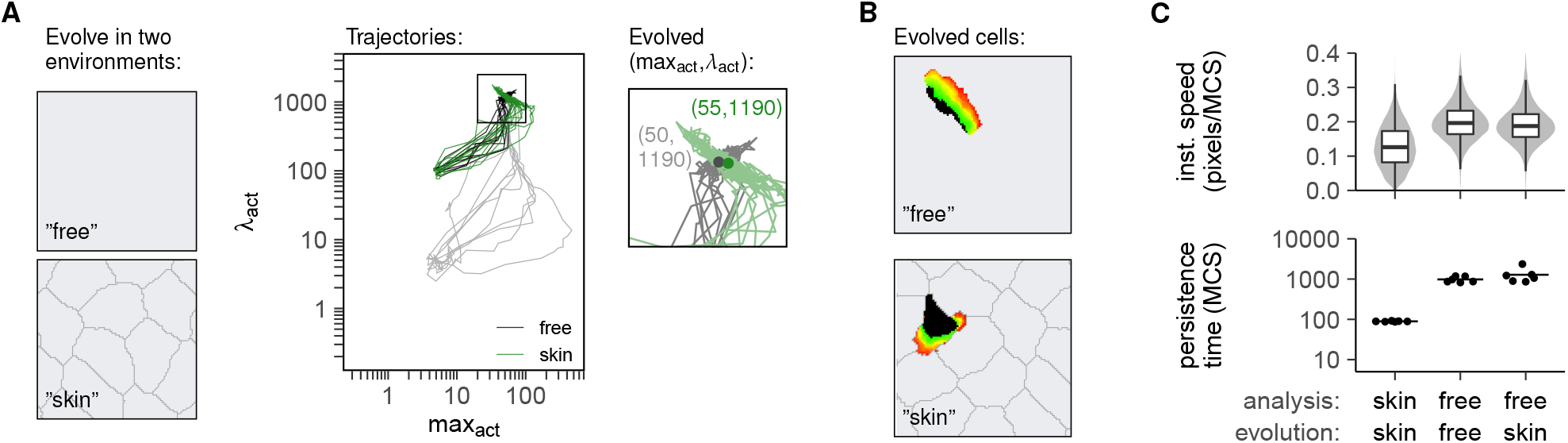
Act cells in different environments evolve similar parameters but different shapes and behaviour. (A) Evolution trajectories of the (max_act_,λ_act_) parameters compared between different runs of evolution in empty space (“free”, see also Figure 2) and evolution in a rigid simulated tissue (“skin”). Black lines represent the “free” cells evolved from the higher λ_act_ = 100, while gray lines show the trajectories from Figure 2. Zoomed square shows where parameters converge in the two different environments after 50 generations, at similar (max_act_,λ_act_) values. (B) Cells evolved in different environments have similar parameters but different shapes and behaviours. See also Movie S1. (C) Speed and persistence of cells with parameters evolved in simulated skin (“skin-skin”), parameters evolved in an empty environment(“free-free”), or parameters evolved in simulated skin but analysed in an empty environment (“free-skin”). Speeds are represented as instantaneous speeds of each individual step in the simulation, and persistence times reflect 6 independent measurements at the same parameters (see Methods for details).

We focus on the epidermal layer of the skin, where skin-resident T cells continuously patrol to search for signs of re-infection by foreign invaders [20]. Because of the skin’s barrier function, the keratinocytes in the epidermis are very tightly packed, forming an extreme example of a restrictive environment. We have previously shown that such restrictions strongly affect cell motion, for example by obscuring the UCSP [13].

Because migration in a stiff tissue requires higher λ_act_ forces [13], we started with a population with slightly higher λ_act_ than before, which was still low enough to prevent active migration (Figure S3A). λ_act_ and max_act_ values once again increased during evolution, and this was associated with an increase in the fitness and the area explored (Figure 4A, Figure S3B,C). All runs once again converged to roughly the same endpoint—this time with max_act_ = 55, λ_act_ = 1190, only slightly different from the endpoint reached by evolving the same cells in a free environment (max_act_ = 50, λ_act_ = 1165, Figure 4A). The small difference was not a result of the higher λ_act_ starting value, because when we used the same starting value for evolution in empty space, cells still stabilised at very similar parameters (max_act_=50, λ_act_=1190, Figure 4A, Figure S3B). Cells in the skin do seem to experience a slightly difference fitness landscape, where the presence of surrounding tissue affects their ability to explore area and stay intact (Figure S3C). Still, the end result was more or less the same: cells evolved towards very similar parameters and once again could *not* reach maximum speed because of increased cell breaking at high λ_act_ values (Figure S3D). Rather, they appear to evolve to a level of speed and persistence that allows them to keep moving within a rigid environment without breaking apart (Movie S1).

Although cells evolved towards remarkably similar parameters regardless of their environment (Figure 4A), they nevertheless differed in shape and behaviour (Figure 4B, Movie S1). Cells evolved in the skin had both a lower speed and persistence than “free” cells in empty space (Figure 4C). To confirm that these differences in migration statistics were not a result of the slight differences in cell-intrinsic parameters, but mostly of the altered environment, we took cells with parameters optimised for migration in the skin (max_act_=55, λ_act_=1190) and analysed their motility in an empty environment. These cells were much more similar to the “free” evolved cells—despite having been optimised in a different environment (Figure 4C, Movie S1). Thus, although cells can optimise their search efficiency to some extent by evolving cell-intrinsic motility parameters, their eventual migration statistics are shaped largely by environmental constraints.

## Discussion

The interpretation of the diversity in T-cell motility patterns as optimised search strategies is remarkably similar to the so-called optimal foraging theory, used by evolutionary biologists to study the ways in which animals search for food [15, 21]. The general idea is that animals (or immune cells) adopt migratory patterns that maximise their ability to find food (or antigens) [34, 35, 22, 25]. Given the implicit assumption that the behaviour we see has somehow been selected during evolution, we seek specific functions the animal (or cell) could have optimised its behaviour for.

Yet even if a certain migration characteristic is optimal or beneficial in some context, this does not prove that it has truly evolved to aid immune system function [36]. It might also have arisen as a side effect of some other process [37], or simply because other migration modes are impossible within the relevant constraints. Ignoring this possibility may lead to spurious interpretations and hamper a true understanding of immune cell migration—especially since it is almost always possible to come up with a context in which the observed pattern would indeed be beneficial [15]. Applying optimality reasoning can thus be seriously misleading when the pattern in question was never really optimised by evolution at all.

We therefore used a CPM to redefine the baseline expectations for T-cell migration behaviour given a realistic intracellular migration machinery, cell shape, and environment. The dynamic interactions in this model yield a complex genotype-phenotype mapping which—unlike random walk models—introduces non-trivial trade-offs and constraints in the speeds and persistences that cells can obtain [13]. This allowed us to use this model to simulate an evolutionary process where we let T cells maximise a very simple objective function: cover as much area as possible. This problem is analogous to destructive search, for which the theoretical optimum behaviour is simply to move as fast and as straight as possible [34]. The evolutionary setting in this model is highly simplified by design, ignoring that T cells may have to optimise different functions simultaneously, might suffer from exploration-exploitation trade-offs [5], and do not evolve individually but as part of a larger organism. Yet even in this very simple setting, T cells do not reach the theoretical optimum because of the constraints and trade-offs naturally arising in the CPM. This problem is exacerbated when cells are placed in a tissue where surrounding cells pose the dominant constraints on T-cell motility: cells with very similar parameters, “optimised” for the exact same function, move in completely different patterns depending on their environment. These results once again highlight that complex migration features can emerge spontaneously from environmental constraints and are not necessarily adapted for some specific function.

Importantly, we also observe migratory behaviours that have previously been attributed to optimality for some T-cell function but are clearly not optimal in our artificial evolution setting. For example, the existence of intervals of fast and slower motion in T-cell tracks has been attributed to an “intermittent search strategy” where T cells balance area exploration (through fast movement) with local exploitation (slower movement and more frequent turning) [5, 25, 6]. Yet we see that variations in speed occur naturally in Act cells: protrusion dynamics automatically yield intervals of slower and faster (or even stop-and-go) motion. In simulated skin, cells move fast when they are moving forward between two keratinocytes, but then must slow down temporarily when they reach a junction and have to choose a direction. Thus, even though cells in our simple evolutionary experiment gain fitness only from exploring area—and not from exploiting it—their motion nevertheless resembles an intermittent search strategy. These observations demonstrate that intermittent search “strategies” can also arise naturally through cell-intrinsic migration dynamics or through the environment, even in cases where they do not benefit any specific function at all.

While the application of optimality theory to animal foraging has provoked considerable criticism and debate [15], the same line of reasoning is applied with far less debate in the context of T-cell migration [5, 38]. Yet our results demonstrate that the criticisms against optimal foraging theory are also relevant for the interpretation of T-cell search patterns. Do T cells in the brain display Lévy-like statistics because that helps them catch rare pathogens [22], or because they are forced to do so by a combination of their cell-intrinsic migration machinery and the structures they are navigating in the brain? Would they adopt a different pattern in the same environment when fighting a more prevalent pathogen, or would they maintain the same migration mode even when it is no longer beneficial? We therefore suggest using models like the CPM [31, 13, 39, 40], where migration patterns arise naturally from an interaction between the cell and its environment rather than being imposed, to define the baseline expectations for T-cell search. By investigating which migratory characteristics emerge without being optimal or even beneficial, we can zoom in on the motility aspects that have truly been evolved to assist immune system function—without being misled by features that are merely inevitable side effects of an intracellular machinery acting in a complex environment.

## Methods

### Act-CPM

For our simulations, we used the Act-CPM [31]. For more information, we refer to the relevant literature [31, 13], but a brief description follows below.

The Act-CPM extends the CPM as described in Figure 1. Every monte carlo step (MCS, the time unit of the CPM), pixels try to “steal” pixels away from neighbouring cells by copying their identity into that pixel. The success probability *P*_copy_ of these copy attempts depend on the *Hamiltonian* (global energy), which consists of different energetic terms:

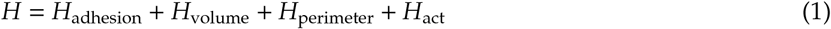

Here, *H*_adhesion_ assigns an energetic penalty to each pair of neighbouring pixels (*i,j*) on the grid that do not belong to the same cell. Likewise, *H*_volume_ and *H*_perimeter_ control the cell’s size and circumference via a penalty that depends quadratically on the deviation from some “target” value 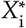 :

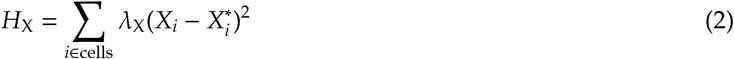

In practice, we mostly look at the energy difference Δ*H* a candidate copy attempt would introduce, rather than considering the absolute energy *H*.

The Act-CPM extends Δ*H* with a positive feedback term, such that pixels newly gained by the cell retain an elevated “protrusive activity” for a period of max_act_ MCS. This is reflected by the negative (=energetically favourable) Δ*H*_act_ assigned to copy attempts that go from a more active source pixel *s* into a less active target pixel *t*:

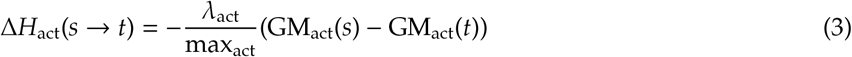

Here, GM_act_(*p*) represents the geometric mean of the activity values in the (Moore) neighbourhood of pixels *p*. Δ*H*_act_ is negative when GM_act_(*s*) > GM_act_(*t*). Details on parameters used will follow below.

### Simulations

All simulations were performed with Artistoo [41]. Simulations were performed for 10,000 MCS during the evolutionary run, or for 50,000 for the detailed simulations to compute speed and persistence (see below).

#### Initialisation

For simulations of “free” T cells moving in an open space, cells were seeded in the middle of a 150×150 pixel grid with periodic boundaries, and allowed a burnin time of 500 MCS to gain their optimal volume and shape.

For simulations of T cells moving in the epidermis, 31 keratinocytes were seeded randomly on a 150×150 pixel grid with periodic boundaries. To ensure proper formation of the tightly packed keratinocyte layer, cells were initially seeded with a tighter perimeter of 200 (making them rounder and preventing cell breaking). Each cell was allowed to grow for 50 MCS before the next cell was seeded, also to ensure that cells did not become entangled and break. After seeding all keratinocytes, the tissue was given 500 more MCS to equilibrate, after which the first keratinocyte was replaced by a T cell and the keratinocytes were given their true perimeter value (see section *CPM parameters* below).

#### CPM parameters

Parameters were selected from [13], allowing realistic shapes and migration behaviour without the cells falling apart (Table 1). Only max_act_ and λ_act_ were varied during the evolutionary runs and in the simulations analysing speed and persistence (see below); other parameters were held constant.

**Table 1:** CPM parameters used in the “free” and “skin” environments. Parameters were kept constant. Skin simulation parameters refer to the T cells; keratinocyte parameters are inside the brackets.

|  | 2D | Skin |
| --- | --- | --- |
| Temperature | 20 | 20 |
| Volume (pixels) | 500 | 500 (750) |
| $\lambda_{\text{Volume}}$ | 30 | 30 (30) |
| Perimeter | 260 | 260 (330) |
| $\lambda_{\text{Perimeter}}$ | 2 | 2 (10) |
| Adhesion cell-background | 20 | 20 (20) |
| Adhesion keratinocyte-keratinocyte | - | 200 |
| Adhesion T cell-keratinocyte | - | 2 |

#### Act-CPM parameters

In the simulations of evolution, λ_act_ and max_act_ were not specified, but evolved spontaneously during the evolutionary run (see section *Evolution of optimal migration modes* below).

To assess speed and persistence at points of interest in the fitness landscapes, simulations were performed at fixed combinations of max_act_ and λ_act_. Along the evolutionary trajectory, simulations were performed at (max_act_, λ_act_) = (5,5), (50,17), (110,30), (70,150), and (50,1185). To examine the behaviour around the optimum, simulations were performed at points surrounding the optimum 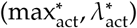 as:

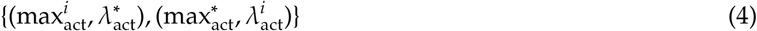

with:

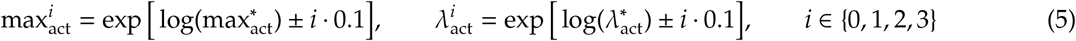

The “optimum” 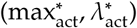 from multiple evolutionary runs was determined as follows. First, runs were removed if they had not “converged” to any optimum (i.e., if their population average of both parameter values changed by ≥20% over the last 10 generations). Parameter values of each remaining run were then averaged over the last 10 generations (focussing only on the 10 fittest individuals and pooling values from different runs) and rounded to the nearest 5. Because populations evolve their parameters on a logarithmic scale (see equation 6 below), this averaging was performed on a log scale as well.

### Evolution

To simulate evolution of optimal migration parameters max_act_ and λ_act_, we used a genetic algorithm as described below. Ten independent runs were performed in every experiment. Simulations in skin were performed in the “stiff” tissue from [13].

#### Evolution of optimal migration modes

To simulate evolution, we started with a population of *N*_pop_ = 10 Act cells. For simulations of cells in an empty environment, cells in the initial population had max_act_ = λ_act_ = 5 (for simulations of cells in the epidermis, initial T cells had the same max_act_ = 5 but a higher λ_act_ = 100 because of resistance from the surrounding tissue). The following steps were then repeated for a total of 50 generations:

1. Population growth: λ = 3 offspring cells were generated from each of the *N*_pop_ cells in the population, with mutated max_act_ and λ_act_ parameters (see section *Mutation* below).
2. Simulation of migration: Each of the (λ + 1)*N*_pop_ cells in the resulting population was simulated independently for 10,000 MCS as described previously, yielding a “fitness” for each cell (see section *Fitness* below).
3. Survival of the fittest: Individuals in the population were ranked according to fitness, and only the *N*_pop_ fittest individuals survived for the next generation.

These choices of *N*_pop_ and λ are somewhat arbitrary; they do not affect our qualitative conclusions on what happens during evolution, but they may affect how long the evolutionary process takes to converge. For example, a larger population allows faster and more thorough exploration of the parameter space. The λ-dependent selection strength determines how long “reasonably fit” individuals can remain in the population and thus have the chance to further evolve. For computational efficiency, we here chose values that allowed evolution to occur within a reasonable number of iterations.

#### Mutation

For mutation of max_act_ and λ_act_ parameters of a given cell, parameter values *x* were first logtransformed and subsequently mutated with a random error term:

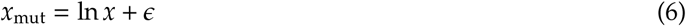

Where

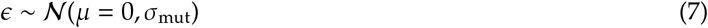

using σ_mut_ = 0.6 for the first 5 simulations and σ_mut_ = 0.2 afterwards. The higher initial choice of σ_mut_ is for efficiency reasons only; note that at the initial low values of λ_act_ and max_act_, cells do not actively move and slight changes in these parameter values can therefore not affect fitness. Only through genetic drift do cells escape this “fitness plateau” into the motile regime where λ_act_ and max_act_ are high enough for migration. Choosing a higher initial σ_mut_ speeds up this process. It does not affect the main results or conclusions from the simulation but simply reduces the number of generations it takes for cells to escape the fitness plateau and start evolving.

#### Fitness

Cells were given a fitness of 0 if they “broke” (connectedness <90% at any point in the 10,000 MCS simulation, see section *Cell breaking*). Otherwise, their fitness equaled the area covered during the simulation (measured in the number of cell volumes of 500 pixels).

### Analysis

In the simulations used to compute speed and persistence, the position of the cell’s centroid was logged every 5 MCS to produce cell tracks. The cell’s integrity was also measured to ensure that cells stayed intact.

Simulated tracks were then analysed in R (version 3.4.4) using the celltrackR package (version 0.3.1) [42]. Speed and persistence were computed in a step-based analysis on 6 groups of 5 simulated tracks (see below), which yielded 6 independent estimates for every parameter combination, from which the mean and SD were assessed. For analysis of mean squared displacements and autocovariance, see below.

#### Speed

To compute speeds, we first computed instantaneous, “step-based” speeds along cell tracks (using the “speed” function of celltrackR). The average of this distribution was then reported as the mean speed.

#### Persistence

The persistence time of moving cells was computed from the decay in the autocovariance curve as described previously in [13].

#### Cell breaking

To quantify cell breaking at a given max_act_ and λ_act_ combination, we counted the percentage of simulations in which the minimum *connectedness* (*C*) was <90%, as described previously [13].

For details, we refer to [13], but briefly: *C* measures the probability that two random pixels of a cell are part of a single, unbroken unit. This measure ensures that an intact cell (which has only *n* = 1 connected component) gets *C*_i_ = 1, whereas a cell broken in many parts (*n* >> 1) gets a very low connectedness. It also means that a single pixel breaking off a cell does not have a huge impact on connectedness, whereas a cell splitting in two equal parts does (even though *n* = 2 in both cases).

#### Mean squared displacement (MSD) curves

Mean squared displacement plots were computed in celltrackR (there are multiple, subtly different methods to compute MSD curves; we used: *aggregate(tracks, squareDisplacement)*). To compare these curves to the persistent random walk model, Fürth’s equation [43, 44] was fitted to these data:

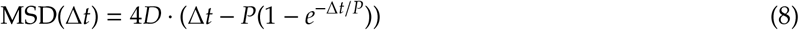

where Δ*t* is the time interval over which displacements are considered, and the persistence *P* and diffusion coefficient *D* are the parameters to be fitted.

To fit these curves robustly, some technical points must be considered. First, there are many more ways to extract small intervals Δ*t* from any given track than there are to extract long ones. In general, if a track contains *n* steps between *t* = 0 and *t* = *n*, there are *n* displacements for Δ*t* = 1 and just one for Δ*t* = *n*. Also note that for larger Δ*t*, many of these overlap: for instance, for Δ*t* = *n* - 1 we have two displacements (*t* = 0 → *n* - 1 and *t* = 1 → *n*), but these overlap almost entirely since they both contain the data between *t* = 1 and *t* = *n* - 1. Thus, they are not independent observations. As Δ*t* increases, we have fewer independent observations of the MSD and thus larger uncertainty in the data. To obtain robust fits, we therefore weighted each data point (Δ*t*, MSD(Δ*t*)) by the number of *independent* displacements the MSD was based on.

Second, CPM cells move on a discrete grid. At timescales far below their persistence time, they only move stochastically—but given the discrete nature of the grid, these very small displacements deviate from the (continuous) Fürth equation and give artefacts when fitting MSD curves. When we are fitting the MSD curve, we are mostly interested in the behaviour around and beyond the persistence time *P*. We therefore fitted curves in two steps:

1. First, a very rough fit was performed on the data. The fitted parameters *D*_0_ and *P*_0_ are not accurate for the reasons mentioned above, but they *are* at least in the right order of magnitude. The estimate *P*_0_ was then used to discard data points with Δ*t* <*P*_0_; thus, we fit only the data at timescales where the cell is actually moving.
2. All points with Δ*t* ≥ *P*_0_ were then used for the final fit using the R function *nls*, setting *“weights”* as described above. Since we are interested in scaling behaviour here and typically consider the MSD on a logarithmic scale, we also perform the fitting on a logarithmic scale:

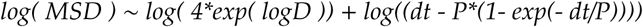

To help the algorithm converge, we fit the logarithm of *D* rather than *D* itself, providing the estimated log *D*_0_ and *P*_0_ to the algorithm as starting point.

#### Autocovariance curves

Autocovariance curves were computed in celltrackR [42] (using: *“aggregate(tracks, overallDot)”*). For persistent random walks, autocovariances should decay exponentially with time interval Δ*t*:

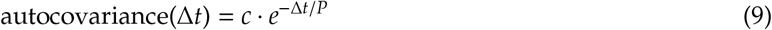

with *c* a constant, and *P* the persistence time.

Once again, we run into the problem that data at very small Δ*t* can cause problems because CPM cells, at that scale, do not actually move (see explanation for MSD curves above). We therefore focused on Δ*t* values that were not too small, filtering Δ*t* > 0.5*P*_MSD_ (with *P*_MSD_ the persistence estimate from the MSD fit).

Given the duration of our simulations, data span a large range of Δ*t* values; however, at low persistences, the autocovariance rapidly decays to zero. If we were to include all the data up to very large Δ*t*, most of these data points would then just contain noise around an autocovariance of ∼ 0 and this noise would dominate the fit. To circumvent this problem, we considered the point *t*_5%_, which is the smallest Δ*t* for which the autocovariance drops below 5% of its initial value. We then filtered points for which Δ*t* < 3*t*_5%_. (The exact choice of this threshold is somewhat arbitrary and does not really matter; the point is that we are looking at a range of Δ*t* values where the autocovariance is actually decaying).

Finally, we fitted the exponential decay equation using R’s *nls* and formula:

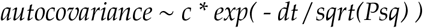

where we fit Psq = *P*^2^ rather than *P* itself to prevent the algorithm from considering negative *P* during the fitting procedure, and provide (*c* = 1, Psq = P_MSD_^2^) as a starting point to help the algorithm converge.

## Code availability

All code required to reproduce this manuscript will be made available on https://github.com/ingewortel/2022-Tcell-evolution upon publication of this manuscript.

## Author Contributions

IW and JT designed the research. IW performed simulations and analysed the data. IW and JT wrote the manuscript.

## Acknowledgements

The authors thank Nir Gov for useful discussions.

## Supplementary Materials

**Movie S1: Tissue constraints impose different migration patterns in evolved cells with very similar parameters**.

Cells migrating in a free environment are broader and move more persistently. This movie is available online at: https://ingewortel.github.io/2022-Tcell-evolution/

**Simulation S1: Area exploration**.

Cellular Potts Model of a migrating Act-cell exploring area. Open this simulation in your browser to see how parameters affect the exploration ”fitness”. Available at: https://ingewortel.github.io/2022-Tcell-evolution/

**Figure S1:**
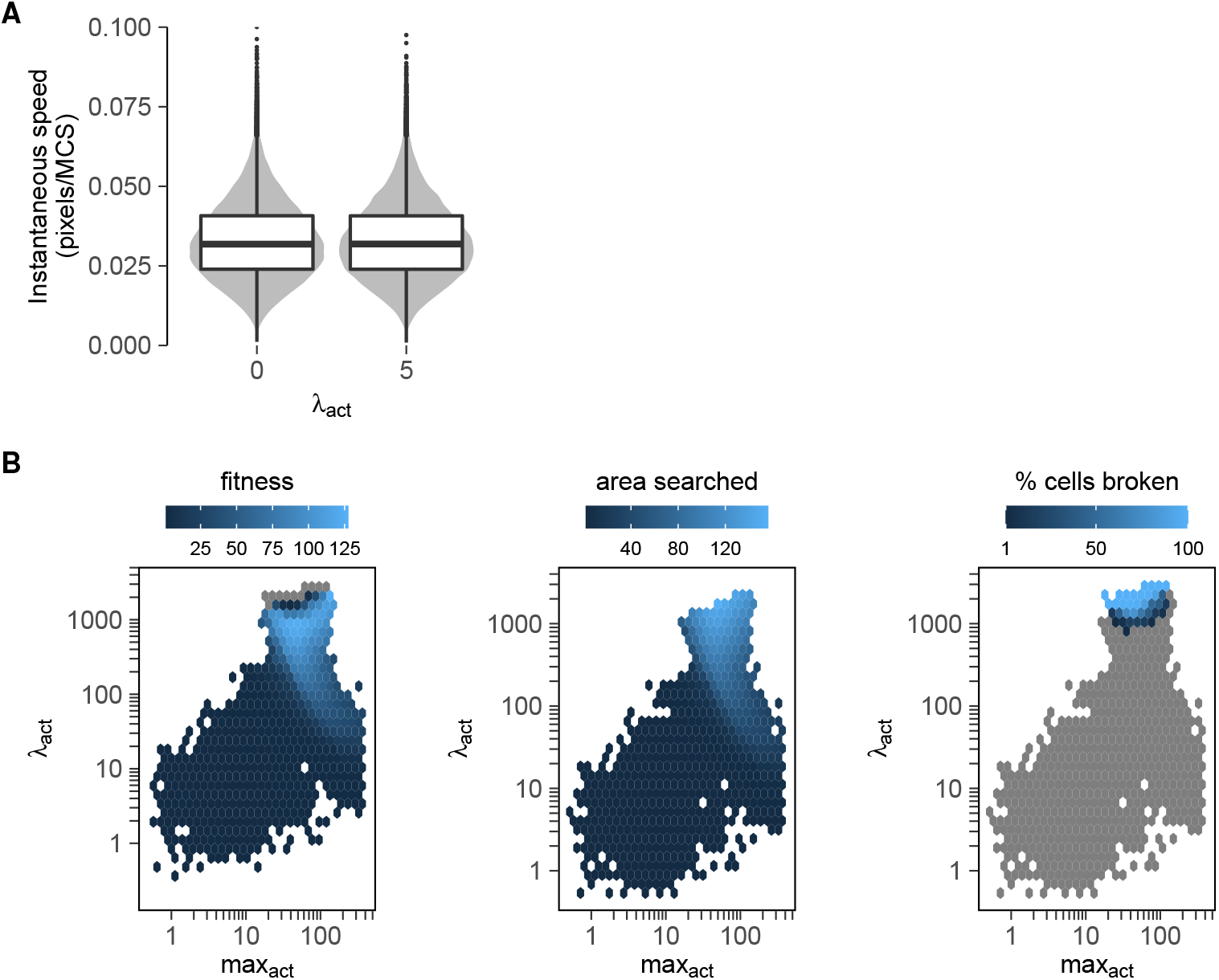
Fitness landscape experienced by cells evolving their max_act_ and λ_act_ values. (A) Act cells with max_act_=5 and λ_act_=5 cannot actively move. Distrubutions of instantaneous speeds equal those of control cells with λ_act_=0 (which cannot form protrusions by definition). (B) Fitness landscape plots show mean fitness (area explored measured in the number of cell target areas of 500 pixels; broken cells have a fitness of zero), mean area searched by non-broken cells, and percentage of broken cells for different (max_act_,λ_act_) combinations. Gray fields represent a value of zero.

**Figure S2:**
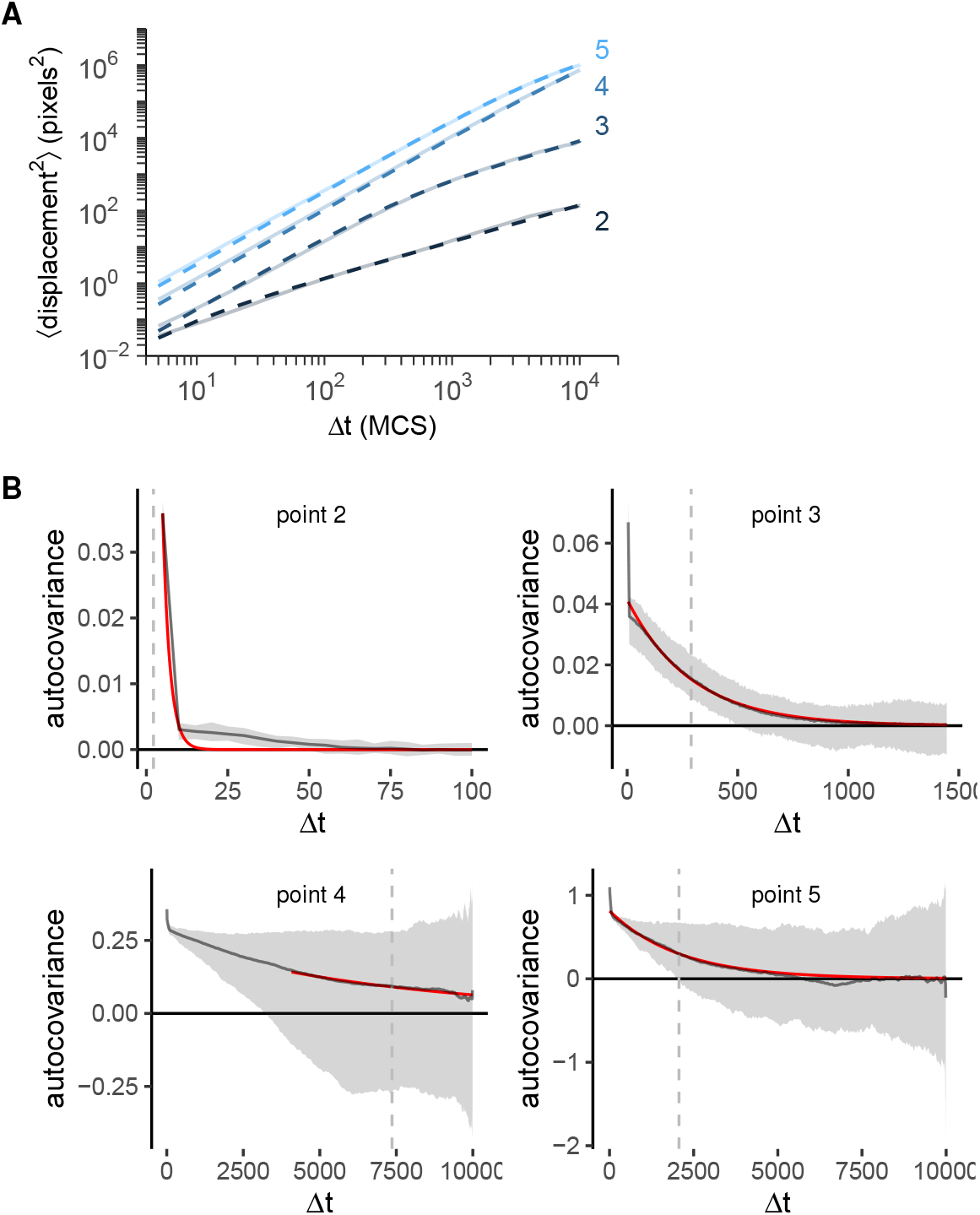
Motion statistics for cells along the evolutionary trajectory. Motion was analysed for the points (2-5) along the evolutionary trajectory of Figure 3 (point 1 was skipped since at these parameters, cells do not yet move). (A) Mean square displacement (MSD) curves of simulated tracks (solid) and the persistent random walk (P-RW) fit (dashed), for points 2-5 along the evolutionary trajectory. (B) Autocovariance curves of the simulated tracks (mean ± interquartile range, gray) and an exponential decay fit (red, autocovariance ∼ exp -Δ*t* /τ). The dashed vertical lines represent the corresponding fitted value of τ, which is another measure of persistence time.

**Figure S3:**
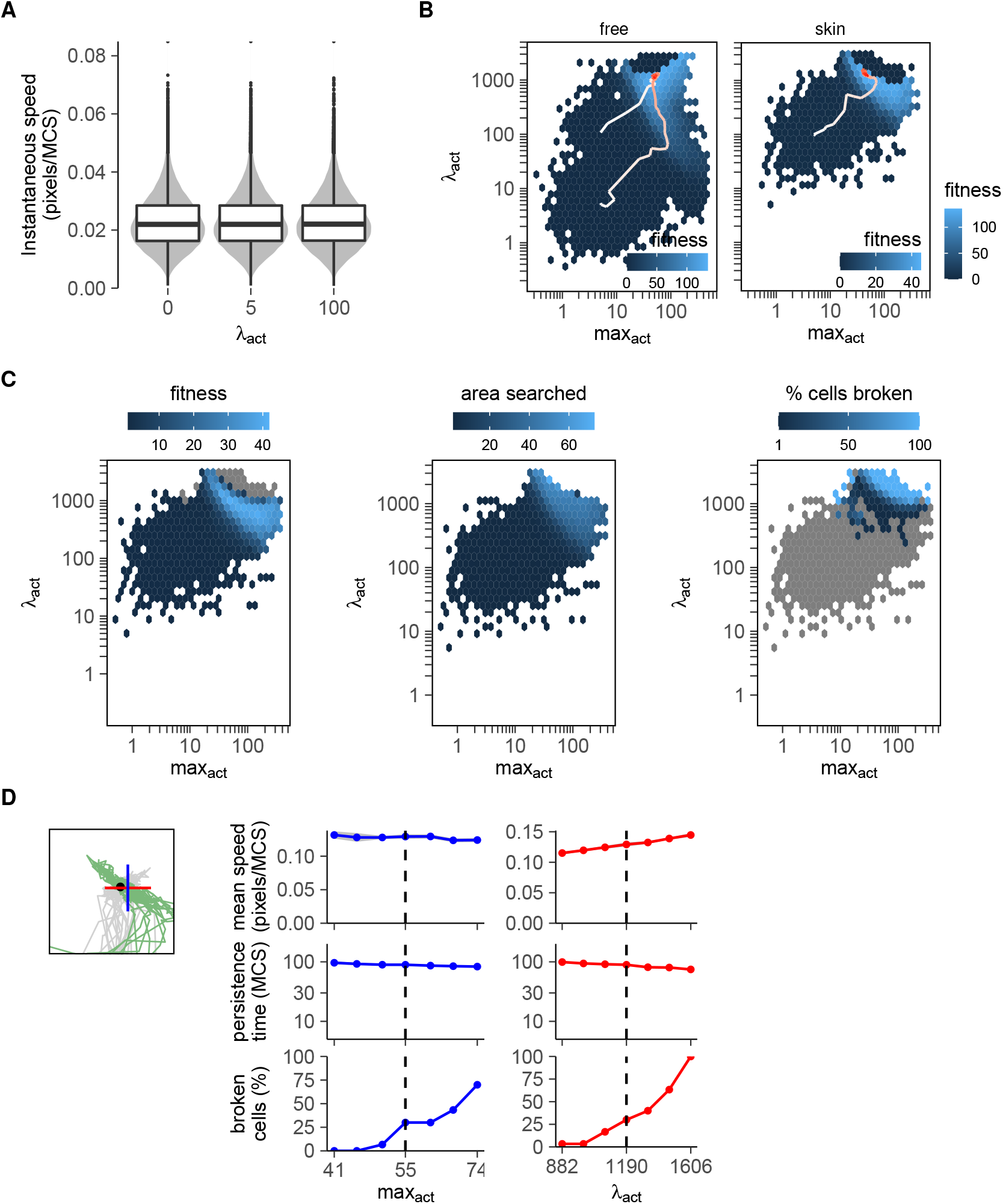
Fitness landscape experienced by cells evolving in a rigid “skin” environment. (A) Cells with max_act_=5 and λ_act_=5 or λ_act_=100 cannot actively move in the rigid skin tissue. Distrubutions of instantaneous speeds equal those of control cells with λ_act_=0 (which cannot form protrusions by definition). (B) Fitness landscape showing median fitness and example trajectories for cells evolved in an empty environment (“free”, two trajectories are shown with a different starting point) compared to cells evolved in stiff tissue (“skin”). (C) Fitness landscape showing mean fitness, mean area searched, and percentage of broken cells (see also Figure S1). (D) Mean speed, persistence, and cell breaking of Act cells in simulated skin at parameters surrounding the evolved optimum (max_act_ = 55, λ_act_ = 1190). The square represents a zoomed version of Figure 4A showing this optimum.

## Footnotes

1 The first paragraph from this introduction also appeared in an earlier preprint [12]. This is because this work started as part of that same project, but it was later decided to split up the project since the work had diverged in two different directions: one focusing on the universal coupling between speed and persistence (now published here [13]), and one focusing on its implications for (evolution of) T-cell search strategies. This introductory paragraph was therefore removed from [13] and maintained in the current work.

